# Axons from the Trigeminal Ganglia are the Earliest Afferent Projections to the Mouse Cerebellum

**DOI:** 10.1101/212076

**Authors:** Hassan Marzban, Maryam Rahimi-Balaei, Richard Hawkes

**Author notes:** Address correspondence to: Dr. H. Marzban, Department of Human Anatomy & Cell Science, Rm 129 BMSB, 745 Bannatyne Avenue, Max Rady College of Medicine, Rady Faculty of Health Sciences, University of Manitoba, Winnipeg, Manitoba, R3E 0J9, Canada **Email:**. **Conflict of Interest:** The authors have no conflicts of interest.

## Abstract

The first stage standard model for the development of afferent pathways to the cerebellum is that ingrowing axons target the embryonic Purkinje cells (E13-E16 in mice). Perinatally and early postnatal (E18-P15) the climbing fibers translocate to the Purkinje cell dendrites, and as the granular layer develops the mossy fibers translocate from the Purkinje cell somata and synapse with granule cell dendrites. In this report we describe a novel earlier stage in the development. Immunostaining for a neurofilament-associated antigen (NAA) reveals the early axon distributions with remarkable clarity. Axons from the trigeminal system enter the cerebellar primordium as early as embryo age (E)9. By using a combination of axon tract tracing, analysis of *neurogenin1* null mice – which do not develop trigeminal ganglia – and mouse embryos maintained *in vitro* – we show that the first axons to innervate the cerebellar primordium are direct projections from the trigeminal ganglia. The data show that the early trigeminal projections are *in situ* before the Purkinje cells are born, and double immunostaining for NAA and markers of the different domains in the cerebellar primordium reveal that they first target the cerebellar nuclear neurons of the nuclear transitory zone (E9-E10), and only later (E10-E11) extend collateral branches to the Purkinje cell plate.

## INTRODUCTION

The cerebellum receives two major afferent types – climbing fibers and mossy fibers. Climbing fibers are exclusively derived from the inferior olivary complex and synapse directly on the Purkinje cell (PC) dendrites. Mossy fiber axons constitute the majority of the afferents. They arise from multiple sources – including the spinal cord and brain stem (reticular formation, vestibular system, trigeminal system etc.) and the cerebrum via the pontine nuclei. The target of cerebellar afferents during embryonic development is the ipsilateral cerebellar primordium, derived from the alar plate of rhombomere 1 (Jacquin et al., 1982, Falls and Alban, 1986, Voogd, 1992, Barmack and Yakhnitsa, 2011, Wingate and Hatten, 1999, Wang and Zoghbi, 2001). The cerebellar primordium houses two distinct germinal zones - the ventricular zone of the 4^th^ ventricle and the rhombic lip (Leto et al., 2016). Fate mapping demonstrates that cells of the 4^th^ ventricle expressing pancreas specific transcription factor 1a (Ptf1+) give rise to all cerebellar inhibitory (GABAergic) neurons (Hoshino et al., 2005), including PCs (delineated by forkhead box P2 (Foxp2) expression:(Fujita and Sugihara, 2012), while the rhombic lip generates the glutamatergic neurons, including granule cells and the large glutamatergic cerebellar nuclear (CN) projection neurons (Fink et al., 2006, Hevner et al., 2006). PCs are born between E10-E13 (all dates are for mice unless stated otherwise): the CN are born slightly earlier (E9-E11) and have a dual origin: the large projection neurons arise from the rhombic lip (Marzban et al., 2014, Leto et al., 2016, Miale and Sidman, 1961, Ben-Arie et al., 1997, Machold and Fishell, 2005), and the nucleo-olivary projection neurons from the ventricular zone (Marzban et al., 2014, Leto et al., 2006, Leto et al., 2008). The products of the germinal zones accumulate in the cerebellar primordium. Based on the expression of transcription factors four distinct domains can be distinguished: c1-c4 (Chizhikov et al., 2006, Zordan et al., 2008). The c1 domain, expressing atonal bHLH transcription factor 1 (Atoh1: (Akazawa et al., 1995, Wang and Zoghbi, 2001)), lies adjacent to the roof plate of rhombomere 1 and comprises cerebellar glutamatergic neurons including the projection neurons of the CN (Fink et al., 2006, Machold and Fishell, 2005, Wang and Zoghbi, 2001). The c2 domain (also known as the Purkinje cell plate) includes Ptf1a+ and Foxp2+ cells and gives rise to all GABAergic neurons, including the PCs. The c3 domain (also known as the nuclear transitory zone - NTZ) is located between c2 and c4 along the entirety of rhombomere 1 except the most rostral end. The glutamatergic CN projection neurons in c3 express LIM homeobox transcription factor 1 alpha (Lmx1a (Chizhikov et al., 2006)) and T box brain gene 1 (Tbr1 (Fink et al., 2006)). Finally, the c4 domain is located rostrally: it is characterized by LIM homeobox protein (Lhx)1/5 expression (Zordan et al., 2008, Chizhikov et al., 2006), but the fates of c4 cells are unknown.

The canonical model of afferent development is that axons enter the embryonic cerebellar primordium between embryo age (E)13-E18 where they terminate on the PC somata (Sotelo and Wassef, 1991, Kalinovsky et al., 2011, Sillitoe, 2016). Climbing fibers enter the primordium between E14 – E15. Postnatally, their synaptic termini translocate from the PC somata to their mature location on the dendritic arbor (Watanabe and Kano, 2011). The earliest mossy fiber projections to the cerebellum are the central processes of the vestibular ganglion neurons (Morris et al., 1988, Ashwell and Zhang, 1998). The cells of the vestibular ganglion are born on E10-E14 (Ruben, 1967) and vestibular afferents from the vestibular nuclei to the cerebellum are first observed at E15 (Morris et al., 1988). Subsequently, as granule cells migrate ventrally from the external granular layer to form the granular layer, the mossy fiber terminals displace from the PCs and synapse with their adult targets, the granule cell dendrites (Marzban et al., 2014, Leto et al., 2016).

In this report we provide evidence for an earlier stage in afferent development. We show that axons of the trigeminal system have already innervated the cerebellum by E9, before the first PCs are born (E10-E13: (Miale and Sidman, 1961)) At this stage, their targets are not the PC somata in c2 but rather the cerebellar nuclear neurons in c3. Subsequently, by E10/11 the first afferents have extended collateral branches to the c2 domain containing the newborn PCs.

## MATERIALS AND METHODS

### Mice

All animal procedures were performed in accordance with institutional regulations and the *Guide to the Care and Use of Experimental Animals* from the Canadian Council for Animal Care. In this study, we used embryos from CD1 mice, and neurogenin1 and neurogenin2 (*Neurog1* and *Neurog2*) null mutant embryos from timed-pregnant mice at E9-E13. Timed-pregnant CD1 mice were obtained from Charles River Laboratories (St. Constant, Quebec, Canada), the University of Calgary Animal Resources Centre, and the Central Animal Care Service, University of Manitoba. Animals were kept at room temperature and relative humidity (18–20°C, 50–60%) on a light and dark cycle (12:12 h) with free access to food and water. The embryo age was determined from the first appearance of a vaginal plug, considered to be E0.5 (all embryo ages in the MS are set to this standard). Mice null for *Neurog1* (Ma et al., 1998) were the generous gift of Dr. Carol Schuurmans (University of Toronto): they were crossed into a CD1 background and compared to normal littermates. Genotyping was performed by polymerase chain reaction as described in (Ma et al., 1998).

### Antibodies

The following antibodies were used:

– Mouse monoclonal anti-NAA (3A10) recognizes a family of neurofilament-associated antigens (NAA:(Bovolenta et al., 1997, Marzban et al., 2008)). Anti-NAA antibody was obtained from the Developmental Studies Hybridoma Bank developed under the auspices of the NICHD and maintained by The University of Iowa, Department of Biological Sciences, Iowa City. It was used diluted 1:1000.
– Rabbit polyclonal anti-ß-tubulin III antibody recognizes neuron-specific class III ß tubulin (Tischfield and Engle, 2010) (Cell Signaling Technology, Danvers Mass.). It was used diluted 1:500.
– Rabbit polyclonal anti-Lmx1a (the gift of M. German, UCSF: used here as a marker of the c3 domain of the cerebellar primordium (Cai et al., 2009)): was generated against a hamster GST-Lmx1a fusion protein. It binds mouse Lmx1a without significant cross-reaction with Lmx1b (Cai et al., 2009) and was used diluted 1:1000.
– Rabbit polyclonal anti-Ptf1a (a marker of the c2 domain of the cerebellar primordium: (Li and Edlund, 2001)) was the gift of H. Edlund (Umeå University, Sweden) and used diluted 1:1000.
– Rabbit polyclonal anti-Forkhead box (Fox)p2 (EMD Millipore, Billarica MA: a marker of PCs in the c2 domain of the cerebellar primordium (Fujita and Sugihara, 2012)) was raised against a GST-tagged recombinant protein corresponding to mouse Foxp2 and used diluted at 1:500.
– Chicken polyclonal anti-T-box Brain Protein 1 antibody (anti-Tbr1: Tbr1 is a neuron-specific T-box transcription factor whose expression during early cerebellar development is restricted to the CN projection neurons of the c3 domain of the cerebellar primordium (Fink et al., 2006)) was raised against a keyhole limpet hemocyanin-conjugated peptide corresponding to mouse Tbr1 (EMD Millipore, Billarica MA) and used diluted 1:500.

### Immunohistochemistry

Pregnant dams were deeply anaesthetized with sodium pentobarbital (100 mg/kg, i.p.) and embryos harvested at the stages of interest and fixed by immersion in 4% paraformaldehyde at 4°C for 48 hours. The embryos were then cryoprotected through a series of buffered 10% (2 h), 20% (2 h), and 30% (overnight) sucrose solutions. Serial series of 20µm thick sagittal or transverse sections were cut through the extent of the cerebellum on a cryostat and collected on slides for immunohistochemistry. Peroxidase immunohistochemistry was carried out as described previously (Bailey et al., 2014). Briefly, tissue sections were washed thoroughly, blocked with 10% normal goat serum (Jackson Immunoresearch Laboratories, West Grove, PA) and then incubated in 0.1M PBS buffer containing 0.1% Triton-X and the primary antibody for 16-18 hours at 4°C followed by a 2 h incubation in horseradish peroxidase-conjugated goat anti-rabbit, anti-chicken or anti-mouse Ig antibodies (diluted 1:200 in PBS; Jackson Immunoresearch Laboratories, West Grove, PA) at room temperature. Diaminobenzidine (DAB, 0.5 mg/ml) was used as the chromogen to reveal antibody binding. Sections were dehydrated through an alcohol series, cleared in xylene and cover-slipped with Entellan mounting medium (BDH Chemicals, Toronto, ON). Embryonic sections for double-label fluorescent immunohistochemistry were processed as described previously (Bailey et al., 2014). Briefly, tissue sections were washed, blocked in PBS containing 10% normal goat serum (Jackson Immunoresearch Laboratories, West Grove, PA) and incubated in both primary antibodies overnight at room temperature, rinsed, and then incubated for 2 hours at room temperature in a mixture of Alexa 546-conjugated goat anti-rabbit Ig and Alexa 488-conjugated goat anti-mouse Ig (Molecular Probes Inc., Eugene, OR), both diluted 1:2000. After several rinses in 0.1M PBS, sections were coverslipped in non-fluorescing mounting medium (Fluorsave Reagent, Calbiochem, La Jolla, CA). Whole mount peroxidase immunocytochemistry was performed according to (Sillitoe and Hawkes, 2002) except that the PBS containing 5% skim milk (Nestlé Foods Inc., North York ON, Canada) plus 0.1% Triton-X 100 (Sigma, St. Louis MO, USA) was used as blocking solution. Biotinylated goat anti-rabbit IgG (Jackson Immuno Research Labs Inc., West Grove PA, USA) was diluted 1:1000 in PBS containing 0.1% Triton X-100 and incubated with the embryos overnight. Embryos were washed with PBS (3 × 2hr) and subsequently incubated overnight with ABC complex solution prepared according to the manufacturer instructions (Vectastain, Vector Laboratories Inc., Burlingame CA., USA). Antibody binding was revealed by using DAB as the chromogen.

### Embryo cultures

Embryo cultures were prepared from E9 CD1 timed-pregnant mice. Each embryo was removed from the amniotic sac and immediately placed in ice-cold Ca^2+^/Mg^2+^-free HBSS containing gentamicin (10 μg/ml) and 6 mM glucose. After positioning the embryos on a membrane (Kimtech Science, Roswell, GA), they were placed into 24-well plates in culture medium supplemented with 10% fetal bovine serum at 100% humidity at 37°C in 5% CO_2_ and maintained *in vitro* for 4 days (DIV).

### Axon tracing

In order to visualize projections from the trigeminal ganglia to the cerebellar primordium dye tracing was carried out according to Ashwell and Zhang, 1998 (Ashwell and Zhang, 1998). Embryos from timed-pregnant dams were immersion-fixed in 4% PFA at 4°C for 2-5 days. One or two crystals of the carbocyanine dye DiI (1,1’-dioctadecyl-3,3,3’,3’-tetramethylindocarbocyanineperchlorate), ~150 μm in diameter, were inserted into the trigeminal ganglion of embryos at ages E10 and E11 (N = 5 each). The trigeminal ganglion was identified by its gross morphology and location, either side of the rostral rhombencephalon and ventral to the angle of the junction between the rostral and the caudal rhombic lips (see asterisk in Fig. 6B). They were then returned to the fixative and stored in the dark at 37°C for up to 8 weeks.

### Imaging and figure preparation

For bright field microscopy, images were captured by using a Zeiss Axio Imager M2 microscope (Zeiss, Toronto, ON, Canada). For low magnification fluorescence microscopy a Zeiss Lumar V12 Fluorescence stereomicroscope (Zeiss, Toronto, ON, Canada) equipped with a camera was used. Images were analyzed by using Zen software. For high magnification fluorescence microscopy, a Zeiss Z1 and Z2 Imager and a Zeiss LSM 700 confocal microscope (Zeiss, Toronto, ON, Canada) equipped with camera and Zen software were used to capture and analyze images. Images were cropped, corrected for brightness and contrast, and assembled into montages using Adobe Photoshop CS5 Version 12.

## RESULTS

### Development of the embryonic cranial nerves

Homogenates of the E11 cerebellum separated by polyacrylamide gel electrophoresis, Western blotted, and probed for NAA immunoreactivity revealed the expected single band of apparent molecular weight ~200kDa (Fig. 1A: (Marzban et al., 2008)). High magnification staining of sections through the cerebellar primordium immunofluorescence stained for NAA indicate that NAA immunoreactivity is localized primarily in axons (Fig. 1B; occasional weakly-immunoreactive somata are seen – not shown). To confirm this interpretation, cerebellar sections were double immunofluorescence stained for NAA and the axon-specific marker β-Tubulin III: the two markers are co-expressed (Fig. 1C-E). (The NAA results were also replicated by using immunostaining for SMI-32 – an alternative anti-neurofilament marker ((Lin et al., 2004): data not shown)).

**Fig. 1.**
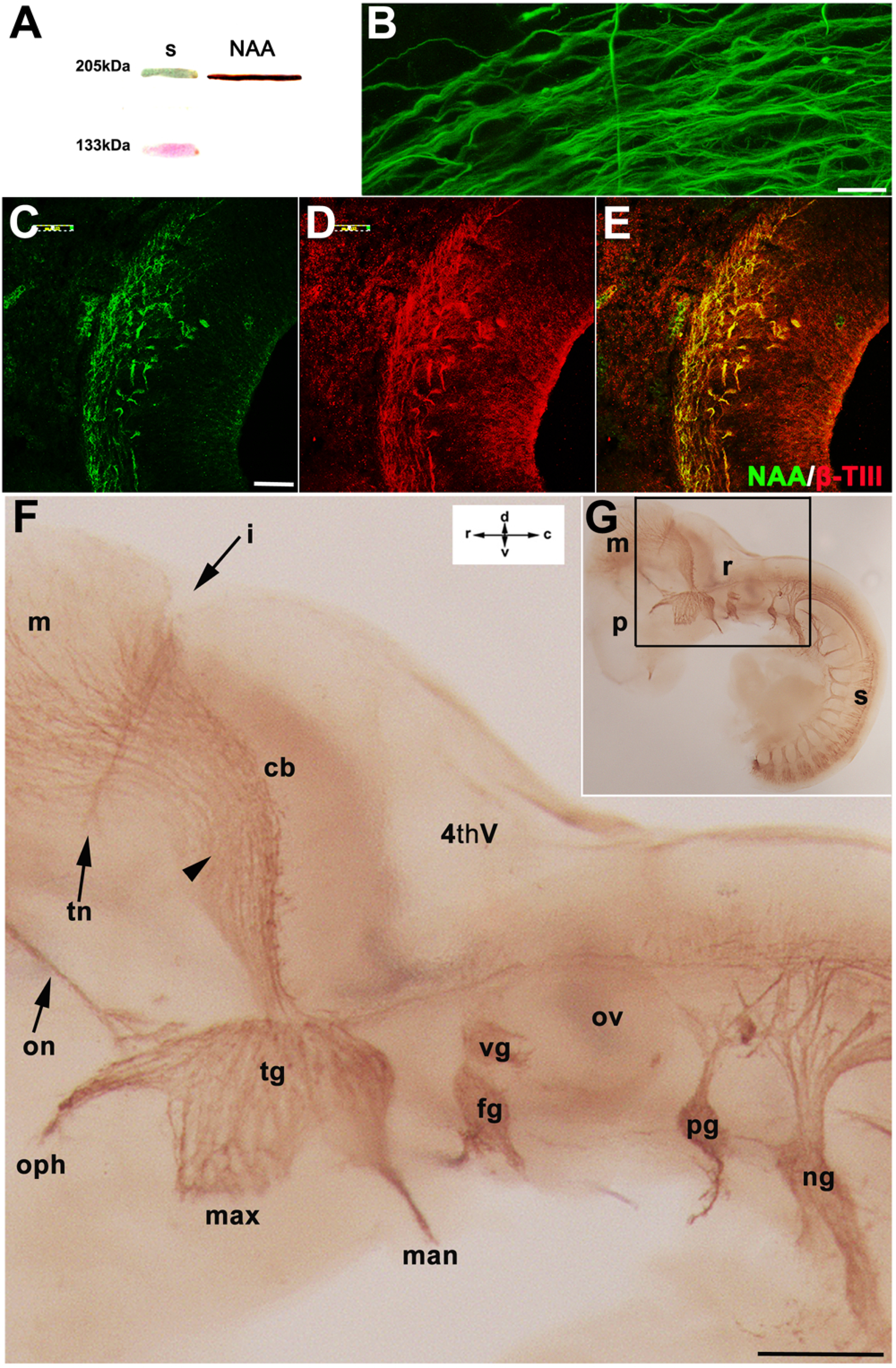
The cranial nerves in the developing mouse embryo are revealed by the expression of neurofilament-associated antigen (NAA). **A**: A Western blots of cerebellar homogenate from the E11 mouse probed with anti-NAA and compared to molecular weight standards (s). The expected single polypeptide band is revealed, with apparent molecular weight ~200 kDa. **B**: High magnification image of sagittal section through the cerebellar primordium at E10. Immunofluorescence staining indicates that the NAA immunoreactivity is associated with axons. **C-E:** A 20µm sagittal section through the E10 cerebellar primordium double immunofluorescence stained for NAA (**E** - green) and ß-tubulin III (ß-T-III: **F –** red; **G** = merged). NAA is co-expressed with ß-tubulin, consistent with an axonal location. **F:** Wholemount anti-NAA immunostaining at E10 from the region outlined in the inset (**G**) shows the developing cranial nerves with a particular focus on relationships between the trigeminal (tg), facial (fg), and vestibular (vg) ganglia and the cerebellar primordium (cb). The trigeminal central projection to the cerebellar primordium is indicated by the arrowhead. The isthmus (i) and the trochlear nerve (tn) indicate the approximate location of the mesencephalic-metencephalic boundary. Additional abbreviations: 4thV = fourth ventricle; fg = facial ganglion (= geniculate ganglion); man = mandibular nerve; max = maxillary nerve; m = mesencephalon; ng = nodose ganglion (= vagus ganglion); on = occulomotor nerve; oph = ophthalmic nerve; ov = otic vesicle; p = prosencephalon; pg = petrosal ganglion; r = rhombencephalon; s = spinal cord. Scale bar: 10 µm in B; 100 µm in C-E; and 200 µm in F

Most cranial nerves originate from the brain stem and have both sensory and motor components. The sensory components, including the ganglia, are born early in development (~E9) and have a dual origin: the distal portions are placodal derivatives and the proximal parts are of neural crest cell origin (Streit, 2004, Schlosser, 2006). To reveal the distribution of NAA immunoreactivity in the embryonic brain, timed embryos were immunoperoxidase stained in whole mount. At E10, NAA immunoreactivity characteristic of phosphorylated neurofilaments is strongly present in both the neuronal somata and axons of the cranial and spinal nerves and ganglia (Fig. 01A,B). In the brain stem, neurons of the epibranchial placodes that comprise the distal parts of the facial (VIIth or geniculate), glossopharyngeal (IXth or petrosal) and vagus (Xth or nodose) cranial nerve ganglia are all NAA+ (Fig. 1A,B). NAA immunoreactivity is also strongly expressed in the trigeminal placode, which lies caudal to the mesencephalon and rostral to the rhombencephalon and forms the large trigeminal ganglion. The peripheral projections of the trigeminal ganglion – the ophthalmic, maxillary and mandibular nerves - follow distinct routes from the outset. The central projections appear to extend rostrally to the cerebellar primordium and mesencephalon and caudally to the rest of the rhombencephalon and the upper cervical spinal cord.

### Axons from the trigeminal nerve are the first to project to the cerebellum

To determine the sequence of cranial nerve development, whole mount and section immunohistochemistry was performed on E9 (Fig. 2) and E11 (Fig. 3) mouse embryos. At E9 NAA immunocytochemistry shows that the trigeminal ganglia and the central projections toward the cerebellar primordium are already present (Fig. 2A, arrow). The peripheral ophthalmic projection is NAA-immunoreactive (Fig. 2A, arrowhead). The facial ganglion has also formed by E9 but the vestibular ganglion, later to be located adjacent to the otic placode, is not seen (Fig. 2C-E: its future location is indicated by the arrow in Fig. 2C: it is present by E10 - see Fig. 1A,B). The centrally-directed axons of the trigeminal ganglion bifurcate upon entry into the brainstem and elongate to form the ascending and descending branches of the trigeminal tract (Fig. 2E) (Erzurumlu and Jhaveri, 1992). At high magnification, the ascending trigeminal axons are seen to terminate in the most rostromedial (Fig. 2E,F) and subpial (Fig. 2E,G) regions of the cerebellar primordium: the c1-c4 divisions are not well defined at E9 but likely this represents the nascent c3. The descending components of the trigeminal fibers are directed caudally (Fig. 2E,H).

**Fig. 2.**
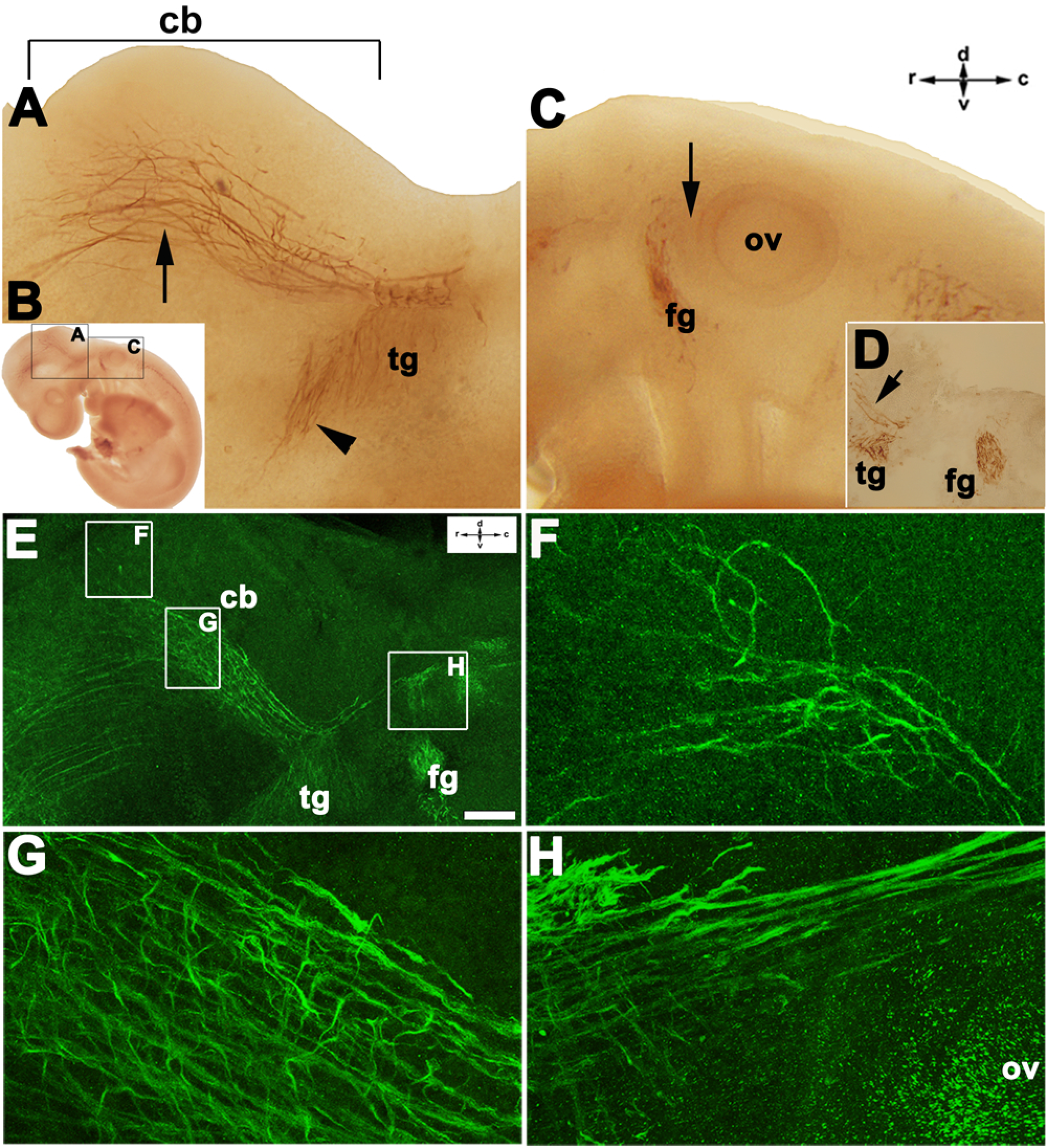
The cranial nerves in the whole mount mouse embryo at E9 immunostained for NAA. **A:** Immunoperoxidase staining of the projection from the trigeminal ganglion (tg) to the cerebellar primordium (cb) at E9, taken from the region indicated in the inset (**B**). The central NAA+ projections – trigeminal or facial or both - extend dorsally to form a bundle (arrow) that continues rostrally to the cerebellar primordium. The ophthalmic peripheral projection from the trigeminal ganglion is indicated by the arrowhead. **C:** Caudal to **A** (the region indicated as **C** in the inset **B**) peroxidase reaction product is deposited in axons of the facial ganglion (fg). The vestibular ganglion has not yet formed (its future location, between the facial ganglion and the otic vesicle (ov), is indicated by the arrow: see also E10 (Fig. 1) and E11 (Fig 3)). **D**: An E9 20µm sagittal section through the rhombencephalon (region indicated as **C** in panel **B**) immunoperoxidase-stained for NAA to show the facial and trigeminal ganglia and the rostral projection (arrow) towards the cerebellar primordium (cb). **E-H**: Confocal microscopy images of whole mount E9 mouse embryos at low magnification (**E**) immunofluorescence labeled for NAA to show the ascending components of the trigeminal ganglion-derived fibers in the rostromedial cerebellar primordium (**F**), the caudolateral cerebellar primordium (**G**), and the descending component of the trigeminal ganglion (**H**). Scale bar: 100 µm in E; 20 µm in F-H

By E11, NAA immunoreactivity clearly reveals the trigeminal, facial, and vestibular ganglia caudal to the cerebellar primordium, together with their peripheral and central projections (Fig. 3A-C). In particular, the central projections of the trigeminal ganglia are directed rostrally towards rhombomere 1 and caudally towards the developing spinal cord (Fig. 3B,C). Caudal to the trigeminal and facial ganglia and adjacent to the hindbrain (rhombomeres 5-6) lies the otic placode - it will give rise to the specialized epithelia of the inner ear, including neurons forming the cochlear–vestibular ganglion (cranial nerve VIII) (Whitfield, 2015, Brown et al., 2003).

**Fig. 3.**
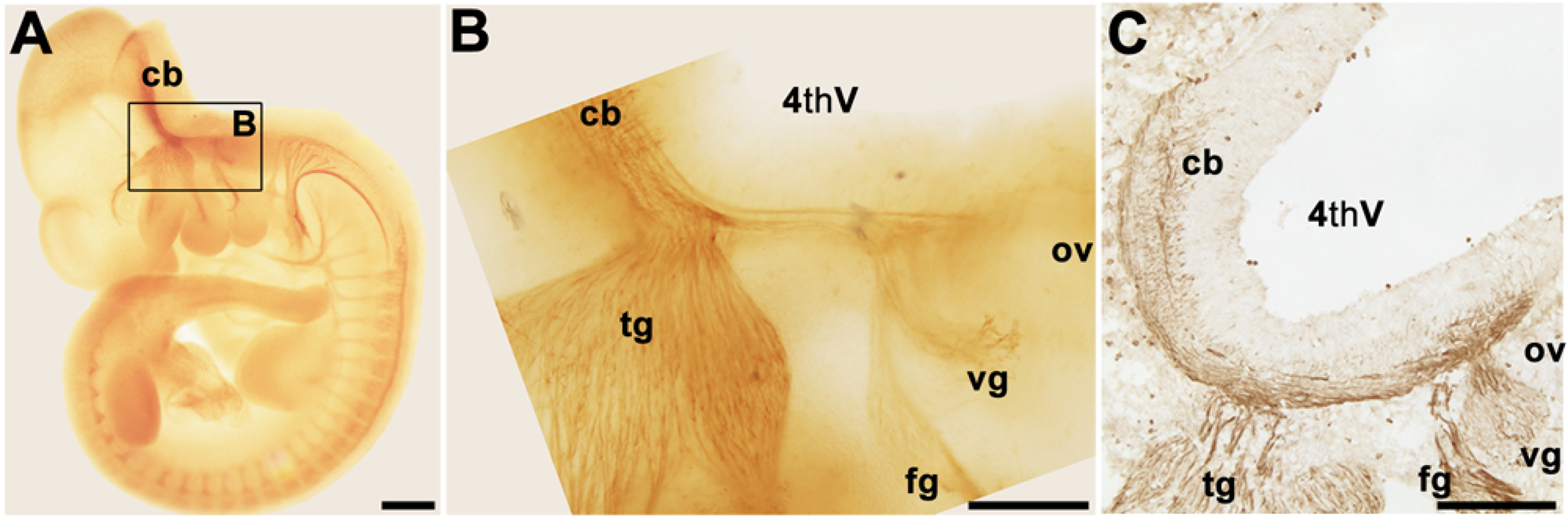
The cranial nerves in the developing mouse embryo at E11 immunoperoxidase-stained in whole mount for NAA. **A:** Low magnification view of an NAA immunoperoxidase stained embryo at E11. **B:** Higher magnification view of the region outlined in “**A”** shows the trigeminal (tg), facial (fg) and vestibular (vg) ganglia. The newly formed vestibular ganglion lies between the facial ganglion and the otic vesicle (ov). An NAA+ axon bundle extends from the ganglia towards the neuroepithelium of the 4^th^ ventricle (4thV) and then rostrally to the cerebellar primordium (cb) and caudally towards the spinal cord. **C**: A 20um sagittal section through the rhombencephalon at E11 immunoperoxidase-stained for NAA. Reaction product is deposited in axons of the developing trigeminal, facial and vestibular ganglia and their central projections to the cerebellar primordium. Scale bars: 500 µm in A; 200 µm in B, C

### Does the trigeminal ganglion project directly to the cerebellum in the embryonic mouse?

To confirm that the embryonic trigeminal ganglion projects directly to the cerebellar primordium at E9-E11, we have pursued three independent strategies – DiI tract tracing, analysis of the *Neurog1* null mouse in which the trigeminal ganglia never form, and analysis of the developing embryo *in vitro*.

First, DiI crystals were applied to the trigeminal ganglia of paraformaldehyde-fixed embryos at E10 and imaged 4-8 weeks later by using confocal microscopy. Care was taken to ensure that diffusing DiI did not impinge upon the nascent vestibular ganglion. Tightly bundled axons were seen pursuing a rostral course across the dorsum of the cerebellar anlage, immediately beneath the pial surface and then descending to enter the cerebellar primordium (Fig. 4A-C). This supports the hypothesis that the trigeminal ganglion projects directly to the embryonic cerebellar primordium and that the axons seen in the primordium at E9 include those from the ipsilateral trigeminal ganglion.

**Fig. 4.**
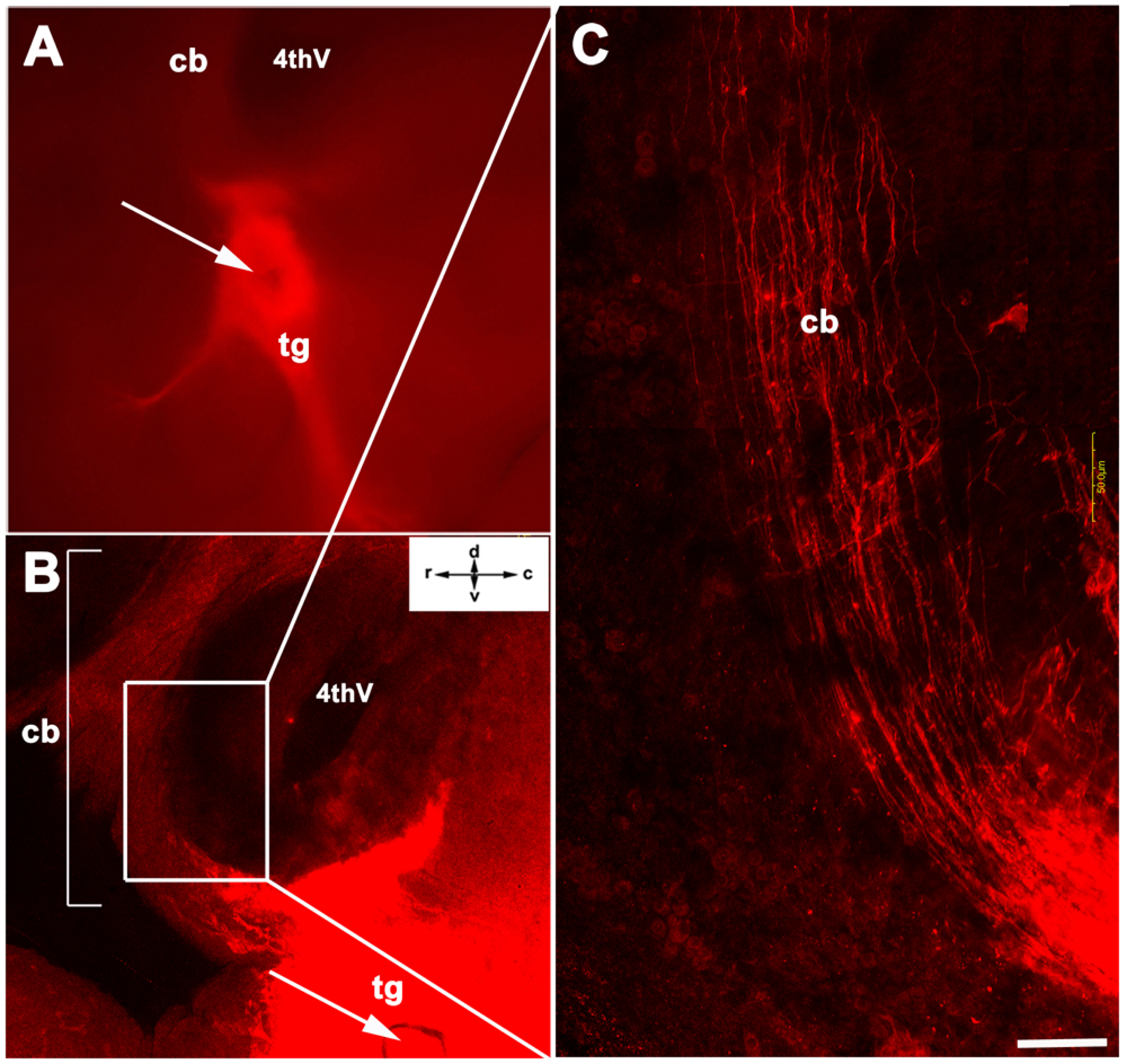
DiI tracing of the projection from the trigeminal ganglion to the cerebellar primordium at E10. **A:** DiI was inserted to the trigeminal ganglion (tg) of a paraformaldehyde-fixed E10 mouse embryo and imaged 3 days later. The arrow indicates the DiI crystal. Axons extend rostrally towards the 4^th^ ventricle (4thV) and the cerebellar primordium (cb). **B:** One month later, the intact embryo was mounted in glycerine, coverslipped and imaged by using confocal microscopy. The arrow shows the location of the DiI crystal in the trigeminal ganglion. The trigeminal ganglion was filled but the DiI did not diffuse as far as the vestibular or facial ganglia. A DiI-stained axon projection is seen extending from the trigeminal ganglion around the 4^th^ ventricle (4thV) into the cerebellar primordium (cb). **C:** A high magnification view of the DiI labeled axon tract between the trigeminal ganglion and the cerebellar primordium showing individual branching axons. Scale bars: 250 µm in A, B; 50 µm in C

Secondly, we wished to confirm that the axon tract between the trigeminal ganglion and the cerebellar primordium derives from the central ganglionic projection and not, for example, the descending projection from the mesencephalic trigeminal nucleus (MesV). The trigeminal ganglion is unique among cranial sensory ganglia in that its neurons are of both placodal and neural crest origin, both of which depend on *Neurog1* function for their development. Indeed it has been shown that elimination of *Neurog1* by homologous recombination prevents the development of the trigeminal and vestibular ganglia (Ma et al., 2000, Ma et al., 1998, Ma et al., 1999, Fode et al., 1998). To confirm that the axons anti-NAA immunostained in the cerebellar primordium at E9 derive from the trigeminal system we investigated the early cerebellar afferent projections in the *Neurog1* null mouse (Fig. 5). Whole mount NAA-immunoperoxidase staining in a control littermate (Fig. 5A,B) and *Neurog1-/-* (Fig. 5C,D) at E9 shows the complete lack of the trigeminal ganglion in the *Neurog1-/-* embryo. Other ganglia are present in their normal locations, albeit somewhat more weakly stained than normal (e.g., the facial ganglion - Fig. 5C,D). The extensive axon ramifications of MesV also appear normal (compare *+/+* in Fig. 5A to the *Neurog-/-* in Fig. 5C). The *Neurog1-/-* cerebellar primordium shows few NAA+ axons at E9 (Fig. 5C,D), while in control littermates the tract is strongly NAA immunoreactive (Fig. 5A,B), consistent with the hypothesis that the tract between the trigeminal ganglion and the cerebellar primordium derives from the trigeminal ganglion rather than from MesV.

**Fig. 5.**
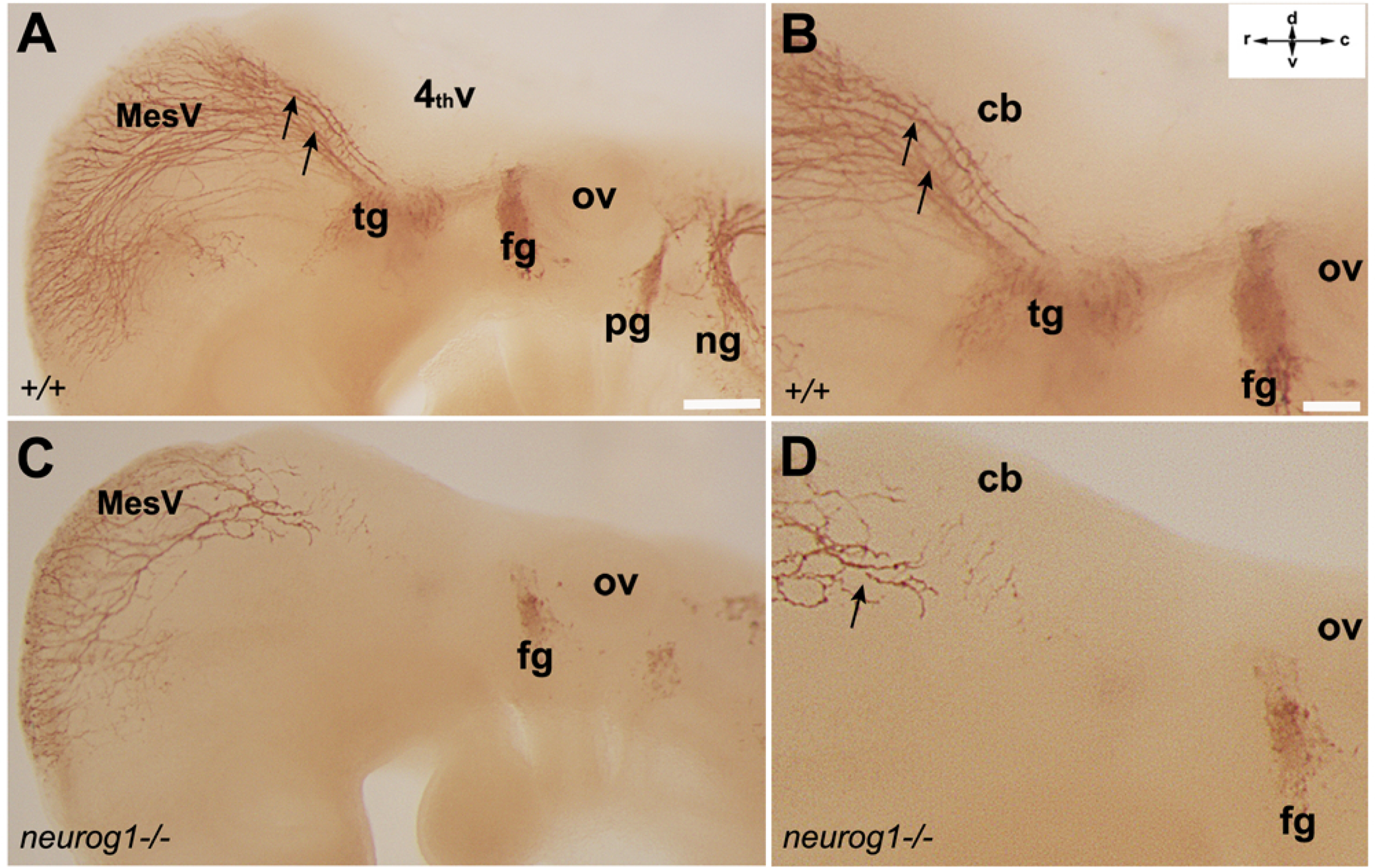
Whole mount NAA-immunoperoxidase staining *Neurog1-/-* and control (*+/+*) littermate embryos at E9. **A:** In the +/+ embryo in the region of the 4^th^ ventricle (4thV) the trigeminal (tg) and facial (fg) ganglia are strongly immunoreactive and NAA-immunoreactive axon projections are seen within the cerebellar primordium (cb: arrow). The vestibular ganglion has not yet developed between the facial ganglion and the otic vesicle (ov). In addition, numerous immunoreactive axons extend caudally from the mesencephalic trigeminal nucleus (MesV) to the cerebellar primordium. **B:** A high magnification view of the embryo in A. **C:** In the *Neurog1-/-* embryo afferents from MesV are present at the rostral end of the cerebellar primordium. The trigeminal and vestibular ganglia are missing and the cerebellar primordium lacks all ascending afferent projections. **D:** A high magnification view of the embryo in **C**. The rostral cerebellar primordium contains a sparse projection from MesV (arrow) but nothing from the caudal cranial nuclei. Scale bars: 200 µm in A, C; 100 µm in B, D

Thirdly, despite the finding that the prominent NAA+ axon tract between the trigeminal ganglia and the cerebellar primordium derives from the central projection of the trigeminal ganglia, NAA immunostaining still indicates the presence of sparse, weakly-immunoreactive axons in the cerebellar primordium of the *Neurog1-/-* embryo (e.g., Fig. 5D - arrow). These might reflect either MesV axons that have already crossed the isthmus at E9 to occupy the rostral end of the cerebellar primordium or perhaps NAA is expressed by the nascent axons of the CN neurons. To test the hypothesis that MesV axons also contribute to the earliest cerebellar afferents, embryos were cultured *in vitro*. In these cultures the axons from MesV completely fail to develop. Embryos were harvested from timed-pregnant CD1 mice at E9.0 and maintained *in vitro* for four days (E9/DIV4: 4 litters - 12 embryo cultures in total). All embryos continued to develop as judged by heartbeat and the development of the neural tube, spinal and cranial nerves etc. (Fig. 6). The overall appearance of the axon tracts in the immunoperoxidase stained E9/DIV4 embryos is not as precise as their age-matched counterparts *in vivo* but most landmarks can be identified and it is clear that significant further development occurred *in vitro* (perhaps to the E11/E12 stage). For example, at E9, the vestibular ganglia have not yet developed (see Fig. 2), but after 4 days *in vitro* (E9/DIV4) they can be identified; the ophthalmic, maxillary and mandibular nerve branches of the trigeminal ganglia can be distinguished; and the trochlear and oculomotor nerves, not yet present at E9 (see Fig. 2A), can be seen after DIV4 (*in vivo* – see Fig. 1F and 3A; *in vitro* – see Fig. 6A,C). The axon tract between the trigeminal ganglion and the cerebellar primordium can be clearly seen (e.g., Fig. 6C,D: arrow). However, the mesencephalon does not develop normally and in particular, the normal descending projections from MesV to the cerebellar primordium are completely absent (compare to the embryos *in vivo* in Fig. 5A,B to the E9/DIV4 embryo in Fig. 6A,C). Despite the absence of NAA-immunoreactive axons from MesV the pathway from the trigeminal ganglion to the cerebellar primordium is well developed (compare Fig. 5A,B to Fig. 6A,C).

**Fig. 6.**
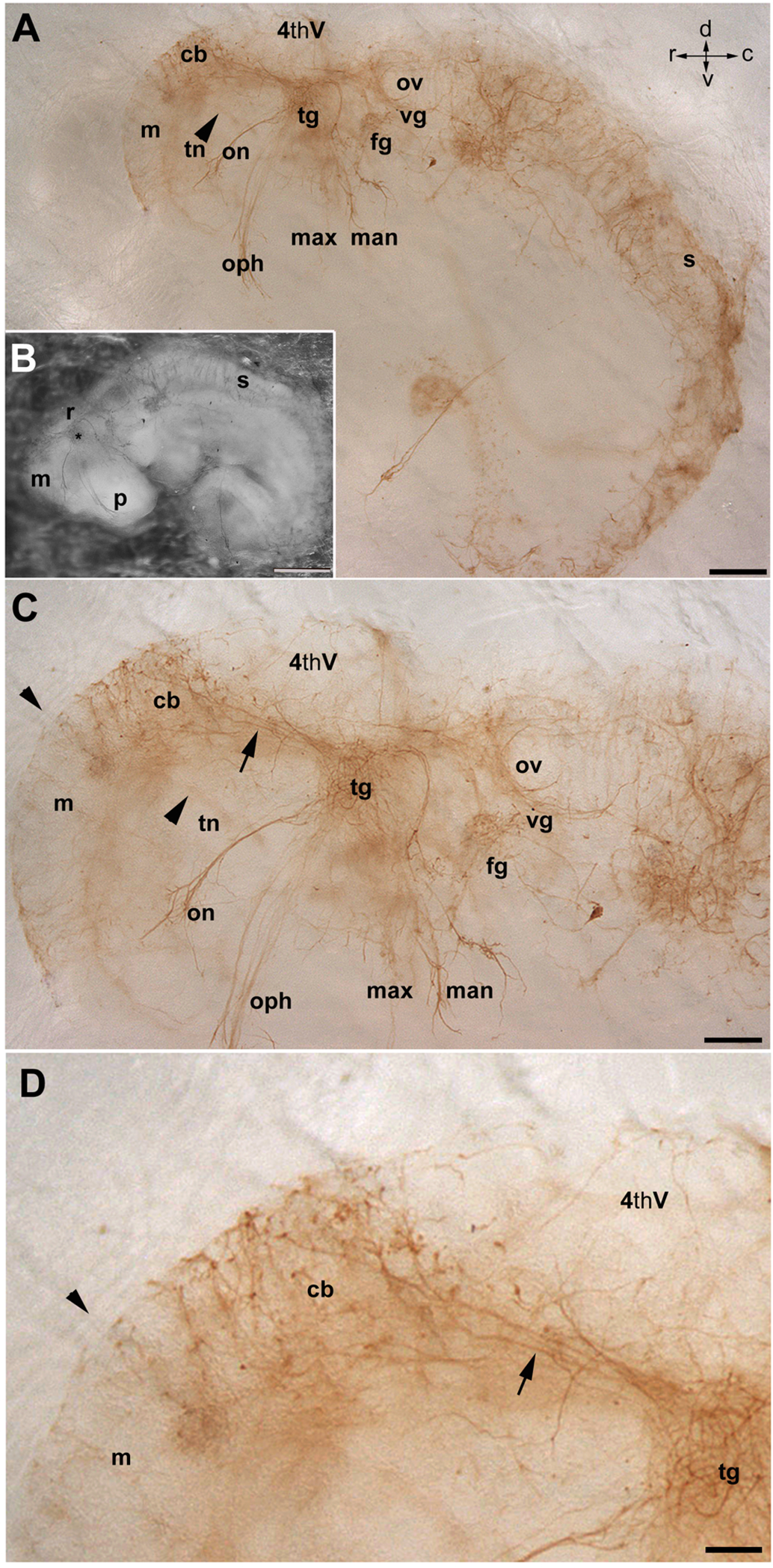
Whole mount culture of a normal embryo taken from a timed-pregnant dam at E9, maintained 4 days *in vitro* (E9/DIV4), and immunoperoxidase stained for NAA. **A:** Anti-NAA immunoperoxidase staining of the E9/DIV4 embryo (shown in bright field in **B**: shows that the spinal and cranial nerves continue to develop *in vitro*. **C:** A higher magnification view of the same E9/DIV4 embryo shows well-developed cranial nerves and the cerebellar primordium. The location of the trochlear nerve (out of the plane of focus in this image) marks the mesencephalic/metencephalic boundary (indicated by an arrowhead). **D:** Higher magnification view of the projection from the trigeminal ganglion to the cerebellar primordium. No NAA immunoreactivity is stained rostral to the mesencephalic/metencephalic boundary (arrowhead: in particular, the MesV is not immunostained – compare to Fig. 5). Abbreviations: 4thV = fourth ventricle; cb = cerebellum; fg = facial ganglion; man = mandibular nerve; max = maxillary nerve; m = mesencephalon; on = oculomotor nerve; oph = ophthalmic nerve; ov = otic vesicle; p = prosencephalon; s = spinal cord; tg = trigeminal ganglion; tn = trochlear nerve; vg = vagus ganglion. Scale bar: 500 µm in A, B; 200 µm in C; 100 µm in D

### The early axons from the trigeminal ganglia terminate on neurons of the developing cerebellar nuclei

Conventionally, it is surmised that the PCs are the targets of the earliest afferents to the cerebellum (Leto et al., 2016). However, the present data show trigeminal ganglion axons have entered the cerebellar primordium by E9, before any PCs are born. This points to a novel early stage in cerebellar circuit development and presumably an alternative afferent target field. To identify the targets of the earliest cerebellar axons, the PC plate (c2) was identified by Foxp2 immunoreactivity (Fig. 7A) and the nuclear transitory zone (c3) by Tbr1 (Fig. 7B) and Lmx1a expression. At E9, double immunofluorescence staining with anti-Lmx1a and anti-NAA shows that NAA+ afferents to the cerebellar primordium are concentrated in the Lmx1a+ c3 domain (Fig. 8A-C). The same distribution is seen at E11 but in addition descending axons are beginning to enter the newly formed c2 (Fig. 8D-F). To confirm that the target of the early afferents to the cerebellar primordium is c3, similar double immunostaining was also performed for NAA and Tbr1. Tbr1 is not expressed in the cerebellar primordium at E9 but by E11 NAA immunopositive fibers are prominent in c3 but not c2 but by E12 NAA+ profiles are plentiful in both c2 and c3 (Fig. 8G-I).

**Fig. 7.**
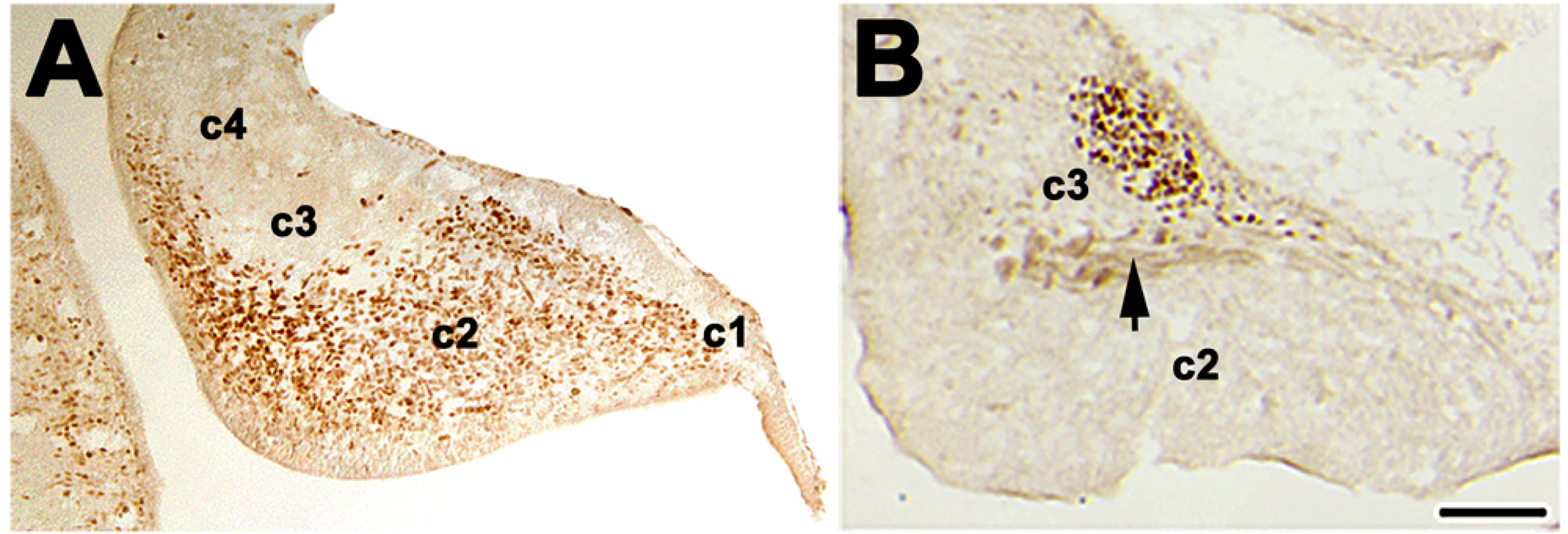
The compartmentation of the embryonic cerebellar primordium. **A:** Anti-Foxp2 immunoperoxidase staining of a 20µm sagittal section through the cerebellar primordium at E13 showing strong immunoreactivity in the cells of c2 (Purkinje cell plate). **B**: Anti-Tbr1 immunoperoxidase staining of a 20µm sagittal section through the cerebellar primordium at E13 shows strong immunoreactivity in c3 (the neurons of the nuclear transitory zone). Weak immunoreactivity is also seen in the afferent projections at the cerebellar core (arrow). Scale bar: 500 µm (A, B).

**Fig. 8.**
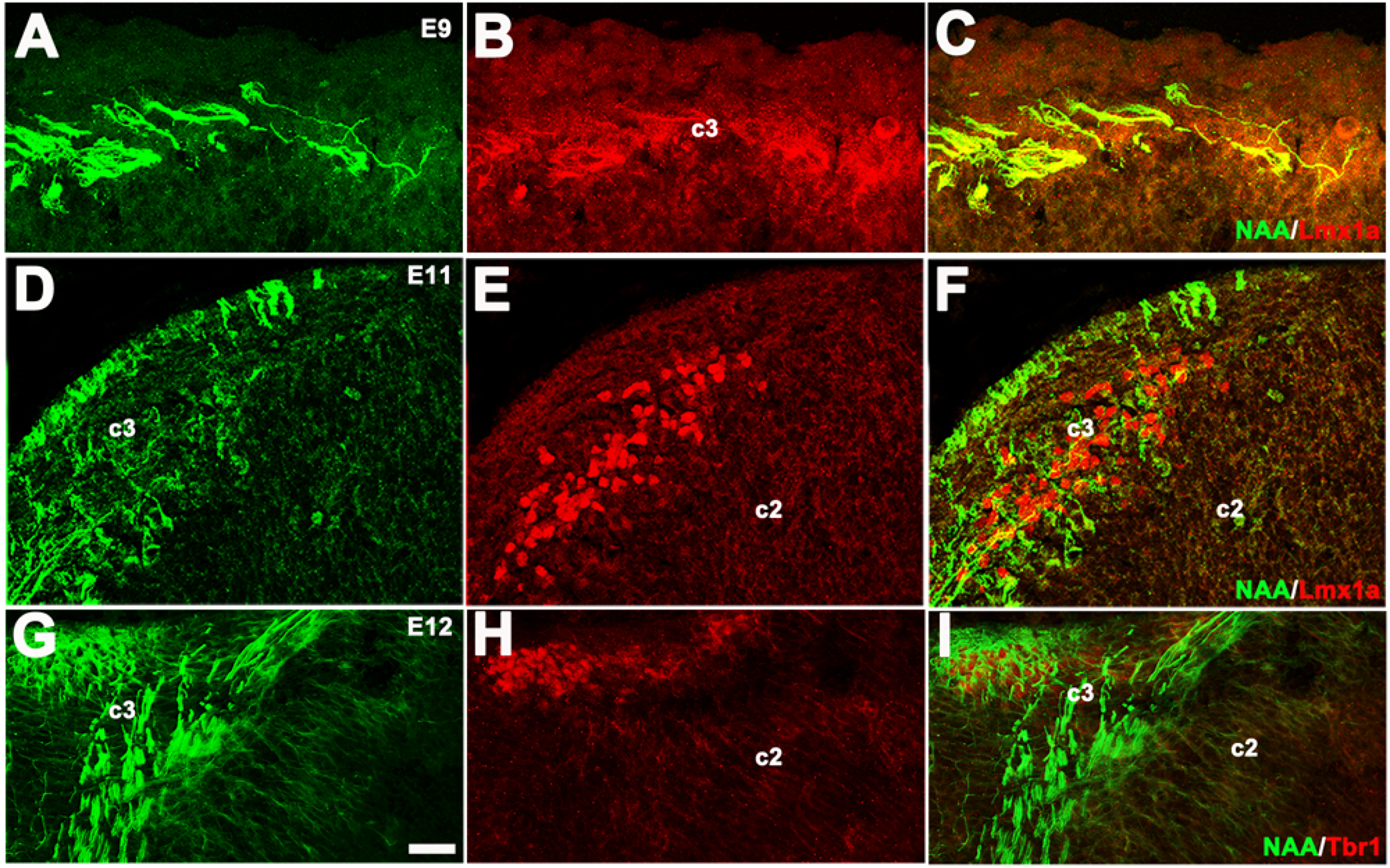
Double immunofluorescence-stained 20µm sagittal sections through the cerebellar primordium to show the development of the early afferent terminal fields. **A-C:** E9 double immunofluorescence staining for NAA (**A** - green) and Lmx1a (**B** - red; **C** - merged) shows that the NAA+ early afferents project to the Lmx1a+ c3 (= nuclear transitory zone). **D-F.** The E11 cerebellar primordium double immunofluorescence stained for NAA (**D** - green) and Lmx1a (**E** - red; **F** - merged). The NAA+ early afferent axons project to the Lmx1a+ cells in c3 but not to the Purkinje cell plate (c2). **G-I**: A sagittal section through the cerebellar primordium at E12 double immunofluorescence stained for NAA (**G** - green) and Tbr1 (**H** - red; **I** - merged) shows the prominent NAA+ afferent projections to c3 (Tbr1+). Scale bars: 20 μm in A-I

The NAA+ axons are not simply passing through c2 but are seen to make intimate contacts with Lmx1a+ neurons with finger-like processes reminiscent of axon growth cones (e.g., Fig. 9). To confirm and explore this further, we exploited two transcription factors expressed by newborn PCs in c2 - Ptf1a and Foxp2 (Hoshino et al., 2005, Fujita and Sugihara, 2012, Hou et al., 2014). As described above, by E11 double immunofluorescence staining of the cerebellar primordium for NAA and Ptf1a (Fig. 10A-C) or NAA and Ptf1a (Fig. 10D-F) shows that NAA+ axons accumulate in c3 and the space between c3 and c2, and then descend into c2. The terminals on the PCs in c2, shown at higher magnification in Fig. 10G and inset, are simple and do not show the extensive envelopment of the cerebellar nuclear neurons characteristic of c3. It is not clear if the projections to c2 and to c3 arise from the same neuronal population (for example, it is possible that the c2 population is the trigeminal ganglion but the later c3 innervation arises from MesV or the vestibular ganglia).

**Fig. 9.**
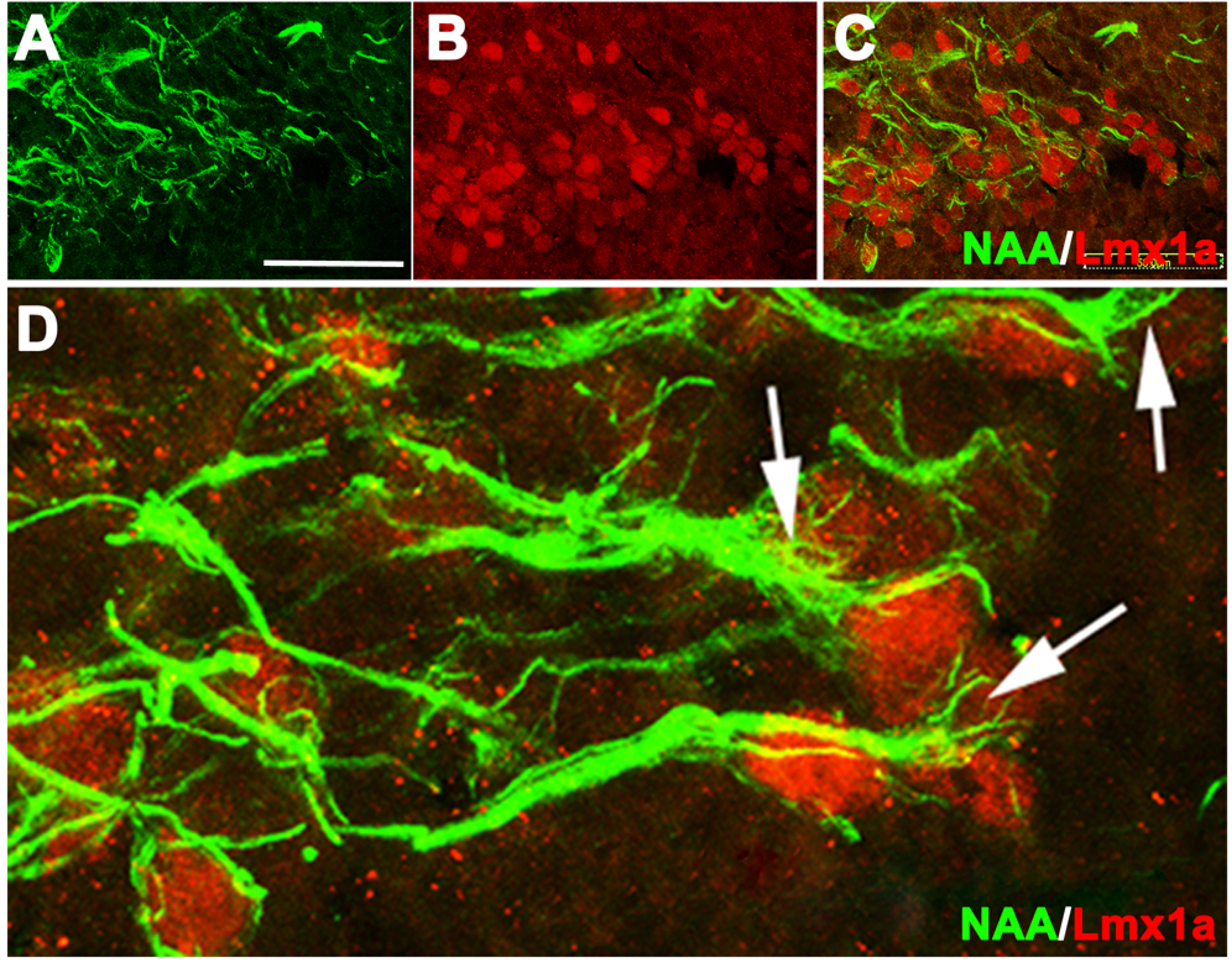
Early afferents in the nuclear transitory zone (c3). **A-C:** 20µm sagittal sections through the E11 cerebellar primordium double immunofluorescence stained for NAA (**A** - green) and Lmx1a (**B –** red; **C** - merged) shows the early afferent axons ramifying throughout c3 (the nuclear transitory zone). **D**: Higher magnification image of c3 double immunofluorescence stained for NAA (green) and Lmx1a (red). Highly branching axon terminals resembling growth cones (arrows) envelop the somata of the Lmx1a+ neurons. Scale bars: 50 μm in A-C

**Fig. 10.**
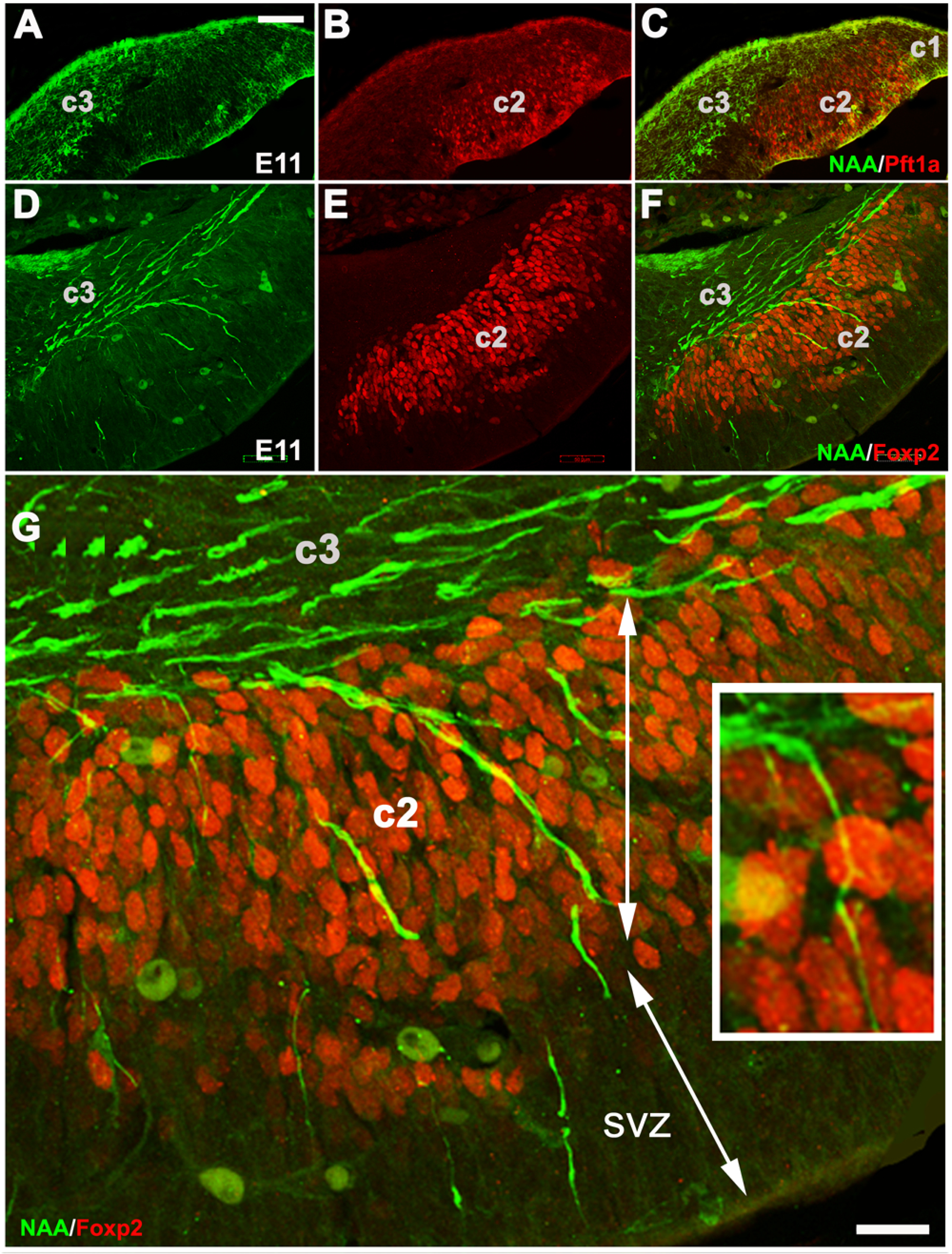
By E11 NAA+ axon projections to the cerebellar primordium are also seen in the Purkinje cell plate (c2). **A-C**: A sagittal section through the cerebellar primordium at E11 double immunofluorescence stained for NAA (**A** - green) and Ptf1a (**B –** red; **C** - merged). Most NAA+ axons are still within c3. (These sections are serial to the NAA- and Lmx1a- immunostained sections in Fig. 8D-F). **D-F**: A sagittal section through the cerebellar primordium at E11 double immunofluorescence stained for NAA (**D** - green) and Foxp2 (**E –** red; **F** - merged) shows axons from c3 extending along the space between c2 and c3 and descending into the Foxp2-immunoreactive c2. **G:** Higher power view of panel **F** to highlight the axons extending from the nuclear transitory zone (c3) into the Purkinje cell plate (c2). In the inset, a single axon is seen – it is slender and unbranched (compare to the terminals in c3 – Fig. 9). In a few cases the axons extend through c2 into the subventricular zone (SVZ). Scattered, large NAA-immunoreactive somata are also seen in c2 – their identity is unknown.

## DISCUSSION

Over a century ago, Tello and Ramon y Cajal (1909) suggested that the earliest afferents to the developing cerebellum originate from the vestibular system and generally studies have focused on the axons of the vestibular ganglia as the earliest source of afferents to the cerebellum: the cells of the vestibular ganglion are born on E10-E14 (Ruben, 1967) and the central processes of the vestibular ganglion neurons enter the cerebellar primordium at E13 (in rat (Ashwell and Zhang, 1998, Morris et al., 1988)). They are not retracted in the adult but become confined to the vestibulocerebellum (e.g., (Carleton and Carpenter, 1984)). In the present study we have identified a novel, earlier stage in the formation of cerebellar afferent topography, which precedes these stages: a direct trigeminal ganglion projection to the cerebellar primordium at E9. This is before the vestibular ganglion has formed (E10-E14 - (Ruben, 1967): see Figs. 1,2) and before the PCs are born. It is also consistent with the observations of Stainier and Gilbert (who suggested that peripheral projections from the trigeminal ganglia emerge as early as E9 (Stainier and Gilbert, 1990)). Because the axon tract in the cerebellar primordium at E9 extends from the trigeminal ganglion via the rhombencephalon and isthmus to the mesencephalon, where MesV is located, it was important to distinguish descending axons from MesV from ascending axons from the trigeminal ganglia. To distinguish between MesV and trigeminal ganglionic projections as the earliest cerebellar afferents, we used a combination of DiI tracing, the *Neurog1* null mouse cerebellum, and embryonic brain cultures. First, DiI tracing supported the proposition that there is a direct axonal connection between the cerebellar primordium and the trigeminal ganglion (Fig. 4). Secondly, we exploited the finding that trigeminal ganglion development is dependent on *Neurog1* function (Ma et al., 1998, Ma et al., 2000)), and elimination of *Neurog1* blocks the development of the trigeminal ganglia (e.g., Fig. 5: (Ma et al., 1998)). The cerebellar primordium in the *Neurog1-/-* mutant lacks trigeminal fibers at E9 (Fig. 5C,D), while the axons of MesV appear normal (Fig. 5C), supporting the hypothesis of a direct trigeminal ganglionic projection to the cerebellar primordium at E9. Thirdly, in cultured embryos NAA+ axons from MesV never develop but the NAA+ tract between the cerebellum and the trigeminal ganglion is nonetheless well developed (Fig. 6). In summary, the data support the hypothesis that the afferent projection to the cerebellar primordium at E9 is an axon tract that originates in the trigeminal ganglion, runs rostrally close to the pial surface, and then targets the nuclear transitory zone (c3).

Within the cerebellar primordium the terminal field of the trigeminal ganglion projection is present at E9 (Fig. 2). This is a day before PCs are first born in the ventricular zone of the 4^th^ ventricle and so at E9 the afferents’ targets cannot be PCs but rather the primary targets of trigeminal ganglion afferents are the neurons of the CN (Fig. 8: CN neurons in c3 are born from E9-E12: (Miale and Sidman, 1961)). It is clear that the innervation of c3 involves intimate interactions with the cerebellar nuclear neurons (e.g., Fig. 9) so they do not appear to be simply axons of passage. Subsequently, NAA+ axons also invade c2 (E10-E11). In theory, these could either be collateral branches of the trigeminal nuclear projection to c2 or a second population of afferents entering the cerebellum along the same pathway but preferentially targeting c2. The absence of NAA+ axons in c2 of the *Neurog1-/-* mouse (Fig. 5) plus the presence of multiple branching axons in the E11 tract (Fig. 4C) favours the interpretation that the c2 innovation derives from collaterals of the same axons that innervate c3, but a novel source cannot be excluded.

Finally, we have avoided calling the direct trigeminal ganglion–cerebellar primordium axons a “mossy fiber” pathway because we are uncertain as whether the tract is preserved in the adult animal or is only present transiently during embryogenesis, in which case a mossy fiber phenotype is moot. In fact, whether the embryonic trigeminal ganglion-cerebellum projection is retained in the adult brain in uncertain. The predominant adult trigeminal projections to the cerebellum arise from the interpolaris and oralis trigeminal nuclei. However, a direct mossy fiber projection from the trigeminal ganglion, via the restiform body to the hemispheres and lateral cerebellar nuclei, has been reported in rat (Jacquin et al., 1982, Marfurt and Rajchert, 1991). It may be that this is the adult version of the embryonic pathway described here, in which case the trigeminal ganglion projection is indeed a mossy fiber pathway (although given that all c2 and c3 receive early afferent input, there must be extensive axon pruning later on). If the sequence of target development – first cerebellar nuclear neurons, then PCs – is replicated by other mossy fiber pathways, this might represent a way by which cerebellar topographical relationships are established so that specific subsets of cerebellar nuclear cells and particular PC stripe subsets are co-innervated to generate the adult modular topography.

## ACKNOWLEDGEMENTS

These studies were supported by grants from the Natural Sciences and Engineering Research Council (HM). We are most grateful to Dr. Carol Schuurmans (U. Toronto) for *Neurog1* null mice, and to Drs. M. German (UCSF) and H. Edlund (Umeå, Sweden) for antibodies.

